# Compensation or preservation? Different roles of functional lateralization in speech perception in older non-musicians and musicians

**DOI:** 10.1101/2023.04.19.537446

**Authors:** Xinhu Jin, Lei Zhang, Guowei Wu, Xiuyi Wang, Yi Du

## Abstract

Musical training can offset age-related decline of speech perception in noisy environments. However, whether functional compensation or functional preservation the older musicians adopt to counteract the adverse effects of aging is unclear yet, so do older non-musicians. Here, we employed the fundamental brain organization feature named functional lateralization, and calculated network-based lateralization indices (LIs) of resting-state functional connectivity (FC) in 23 older musicians (OM), 23 older non-musicians (ONM), and 24 young non-musicians (YNM). OM outperformed ONM and almost equalized YNM in speech-in-noise/speech tasks. In parallel, ONM exhibited reduced lateralization than YNM in LI of intrahemispheric FC (LI_intra) in cingulo-opercular network (CON) and interhemispheric heterotopic FC (LI_he) in language network (LAN). Moreover, OM showed higher neural alignment to YNM (i.e., similar lateralization pattern) than ONM in LI_intra in CON, LAN, frontoparietal network (FPN) and default mode network (DMN) and LI_he in DMN. These findings suggest that musical training contributes to the preservation of youth-like lateralization in older adults. Furthermore, stronger left-lateralized and lower alignment-to-young of LI_intra in somatomotor network (SMN) and dorsal attention network (DAN) and LI_he in DMN correlated with better speech performance in ONM. In contrast, stronger right-lateralized LI_intra in FPN and DAN and higher alignment-to-young of LI_he in LAN correlated with better performance in OM. Thus, functional preservation and compensation of lateralization may play different roles in speech perception in noise for the elderly with and without musical expertise, respectively. Our findings provide insight into successful aging theories from the unique perspective of functional lateralization and speech perception.

**Significance statement:** As a positive lifestyle which contributes to neural resource enrichment, musical training experience may mitigate age-related decline in speech perception in noise through both functional compensation and preservation. What is unknown is whether older musicians rely more on one of these mechanisms, and how is it different from older non-musicians. From a unique perspective of functional lateralization, we found that high-performing older musicians showed stronger preservation of youth-like lateralization with a more right-lateralized pattern whereas high-performing older non-musicians were associated with stronger scaffolding of compensatory networks with a more left-lateralized pattern. Our findings suggest that older musicians and non-musicians exhibit different coping strategies in terms of functional lateralization against aging, which would largely enrich aging theories and inspire training intervention.

## Introduction

The world’s population ages at an unprecedented rate, posing a serious burden on families and society. Neurocognitive aging is characterized by multidimensional cognitive decline and prominent changes in brain structure and function (1–3). One of the salient problems for older adults in everyday life is difficulty understanding speech in “cocktail party” scenarios, even if their hearing is normal for their age. Both the hemispheric asymmetry reduction in older adults (HAROLD) model and the scaffolding theory of aging and cognition (STAC) posit that our brain could adapt through compensatory mechanisms to offset the adverse effects of aging, including increased fronto-parietal and bilateral recruitment with decreased brain lateralization (1, 4). Meanwhile, based on the revised STAC model (5), life-course factors such as musical training can enhance both the functional preservation and compensatory scaffolding processes to ameliorate the aging effects. Indeed, empirical studies have demonstrated the benefits of long-term or short-term musical training on speech perception in noisy environments for older adults (6–10), although such benefits have not been robustly observed in young and middle-aged adults (11, 12). However, it remains unclear whether and how functional preservation and compensation differentially support speech in noise perception in older adults with or without lifetime musical training experience. The answer to this question could help us develop effective and individualized interventions to promote listening skills and healthy aging in the elderly.

On the one hand, older adults may engage in functional compensation to improve speech perception in noisy environments. This compensation is achieved through elevated activities in the cingulo-opercular network (CON, also called salience network) and frontoparietal network (FPN), supporting a compensatory scaffolding for attention, working memory and cognitive control (13–16). Additionally, old adults also up-regulate task activity and preserve the specificity of phoneme representations in sensorimotor regions to compensate for speech perception in noise (17). These functional compensations result in a reduction of the asymmetry of activation patterns associated with aging, as proposed by the HAROLD model (3, 4), and are supported by behavioral (18), positron emission tomography (PET) (19), electroencephalography (20), near-infrared spectroscopy (21), as well as task and resting-state functional magnetic resonance imaging (fMRI) studies (22, 23). These findings support the compensation theory, which proposes that brain networks become more bilateral to compensate for age-related neural declines.

Older musicians may also employ functional compensation to mitigate the aging effect on speech perception. Compared to non-musicians, musicians exhibit enhanced auditory-motor integration during speech perception in noise, manifested by increased right lateralization of the dorsal stream’s key fiber (24), stronger activation of the right auditory cortex, and heightened functional connectivity between the right auditory and bilateral motor regions (25). Actually, greater musical training-related plasticity were reported in the right hemisphere at both the structural and functional levels (26). A recent study has shown the increased recruitment of frontal-parietal regions and greater deactivation of the default mode network (DMN) region as two means of functional compensation for speech in noise perception in older musicians (10)(Zhang et al., 2023).

On the other hand, musical training has been suggested as a “cognitive reserve” that can delay age-related cognitive declines (6, 7). Compared to older non-musicians, older musicians show enhanced central auditory processing functions and preserved cognitive abilities including auditory attention and working memory, which may support their comparable performance to young adults (8, 9, 27, 28). Moreover, musical expertise in young adults has been found to retain right-lateralized ventral attention and improve the neural specificity of speech representations in auditory as well as speech motor regions (25, 29), whereas maintained youth-like neural specificity of speech representations in sensorimotor areas in older musicians has been revealed as a key mechanism to achieve successful speech in noise perception (10). Therefore, the benefit of musical expertise on older adults’ speech perception in noisy environments is likely associated with functional preservation as well.

In this resting-state fMRI study, we aimed to reveal the specific brain mechanism that older musicians (OM) and older non-musicians (ONM) use to counteract age-related decline, by uncovering how aging and life-long musical training experience affect intrinsic functional lateralization and its relationship with speech in noise perception ability. We adopted functional lateralization based on spontaneous brain activity at resting state, as it reflects a fundamental organization characteristic of the intrinsic functional architecture of the brain and changes with aging and musical experience (23, 30, 31). Previous studies have shown that the degree of functional lateralization of resting-state functional connectivity (rs-FC) predicts individuals’ language and visuospatial abilities, providing evidence that functional lateralization of rs-FC is associated with human cognition (32–34). Here, we defined two types of lateralization indices (LIs) based on rs-FC and compared them between older groups and young non-musicians (YNM): LI of intrahemispheric FC (LI_intra) which represents the left-right connectivity strength difference within the same hemisphere, and LI of interhemispheric heterotopic FC (LI_he) which represents the left-right connectivity strength difference across the bilateral hemispheres (30, 33–35). Larger positive values of LI_intra and LI_he indicate stronger within-hemisphere interactions and across-hemisphere interactions in the left hemisphere, respectively, whereas larger negative values imply stronger interactions in the right hemisphere. To further figure out the functional lateralization of which network shows functional compensation and which one produces functional preservation in the aging process with and without musical expertise, we performed correlations between speech in noise perception threshold and network-based LI as well as its neural alignment-to-young, which was defined by inter-subject spatial pattern similarity between LI of each older subject and group average of LI in YNM across the same network vertices (see **Fig. 1**). We hypothesized that ONM and OM would adopt different coping strategies against aging, by showing different functional lateralization patterns and relationships between network-based LI and/or its neural alignment and speech perception thresholds. Specifically, ONM may exhibit functional compensation, that is, the less similar the lateralization was to YNM, the better performance. And OM were more likely to reflect functional preservation, that is, the more similar the lateralization was to YNM, the better performance.

**Fig. 1.**
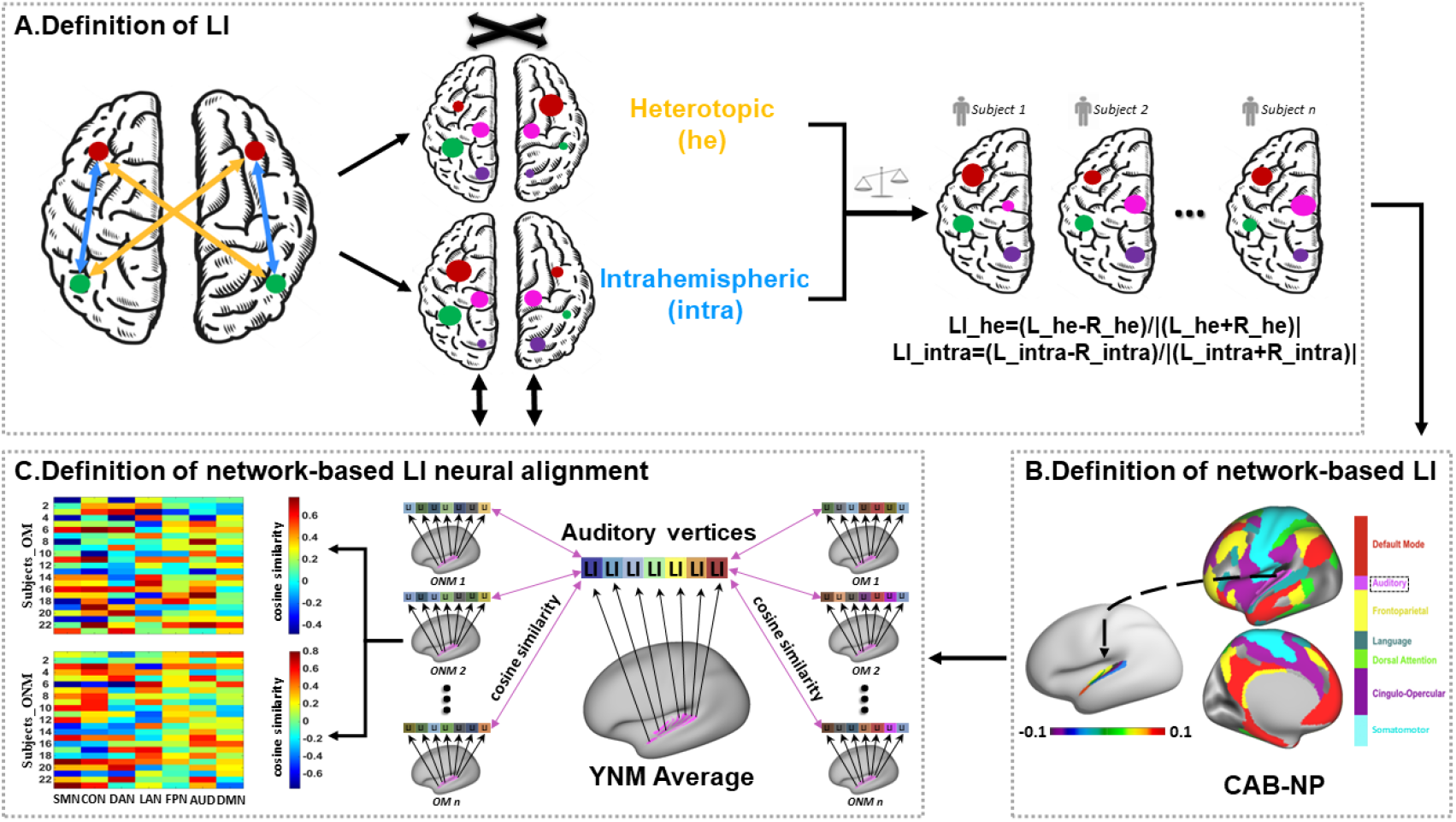
Workflow of analyses. (**A**) Definition of LI. We first defined two different types of functional connectivity (FC) among the whole brain, named interhemispheric heterotopic FC (yellow) and intrahemispheric FC (blue). For a specific surface vertex, the heterotopic (he) was defined as the sum of heterotopic FCs between this vertex and all the others in the opposite hemisphere except the homotopic one, whereas the intrahemispheric (intra) was defined as the sum of intrahemispheric FCs between this vertex and all the others within the same hemisphere. Then, the functional lateralization between each homotopic pair of surface vertices was quantified by a commonly used laterality index (LI): LI = (L-R)/|(L+R)|. Therefore, every subject would have two LI maps of he and intra. Larger positive values of LI_he and LI_intra imply stronger across-hemisphere interactions or within-hemisphere interactions in left-hemispheric vertices, respectively, whereas larger negative values indicate stronger interactions in right-hemispheric vertices. (**B**) Definition of network-based LI. Based on Cole-Anticevic Brain-wide Network Partition version 1.0 (CAB-NP v1.0), we acquired LIs of homotopic vertices belonging to seven task-relevant networks including somatomotor (SMN), cingulo-opercular (CON), dorsal attention (DAN), language (LAN), frontoparietal (FPN), auditory (AUD), and default-mode (DMN). (**C**) Definition of network-based LI neural alignment. Taken the AUD for example, we calculated cosine similarity between the group average of LI in young non-musicians (YNM) and LI in every older subject across all AUD vertices. By doing this, two cosine similarity matrices for seven networks of older non-musicians (ONM) and older musicians (OM) were obtained. Using the same approach, we also obtained the cosine similarity matrix of YNM.

## Results

### Musical expertise counteracts age-related decline of speech perception in noisy environments

Twenty-four YNM, 23 OM, and 23 ONM completed the experiment. The two older groups exhibited no significant differences in age, hearing level and Montreal Cognitive Assessment (MoCA), but significant difference in education (*t*_44_ = 3.769, Cohen‘s *d* = 1.112, *p* < 0.001). Among three groups, separate one-way analysis of variance (ANOVA) showed a significant main effect of group on speech perception threshold with noise masking (speech-in-noise (SIN): Welch *F*(2, 38.091) = 25.070, *η*^2^ = 0.526, *p* < 0.001), speech perception threshold with speech masking (speech-in-speech (SIS): Welch *F*(2, 38.920) = 28.217, *η*^2^ = 0.568, *p* < 0.001), and auditory digit span (*F*(2, 67) = 28.749, *η*^2^ = 0.462, *p* < 0.001). Post-hoc analysis revealed that ONM showed significantly worse performance on each task than OM and YNM [all false discovery rate (FDR)-or Games-Howell-corrected *p* < 0.001], but no significant difference was found between OM and YNM, suggesting that musical expertise offset older adults’ decline of speech perception in noisy environments. For more details, see **Table 1**.

**Table 1.**
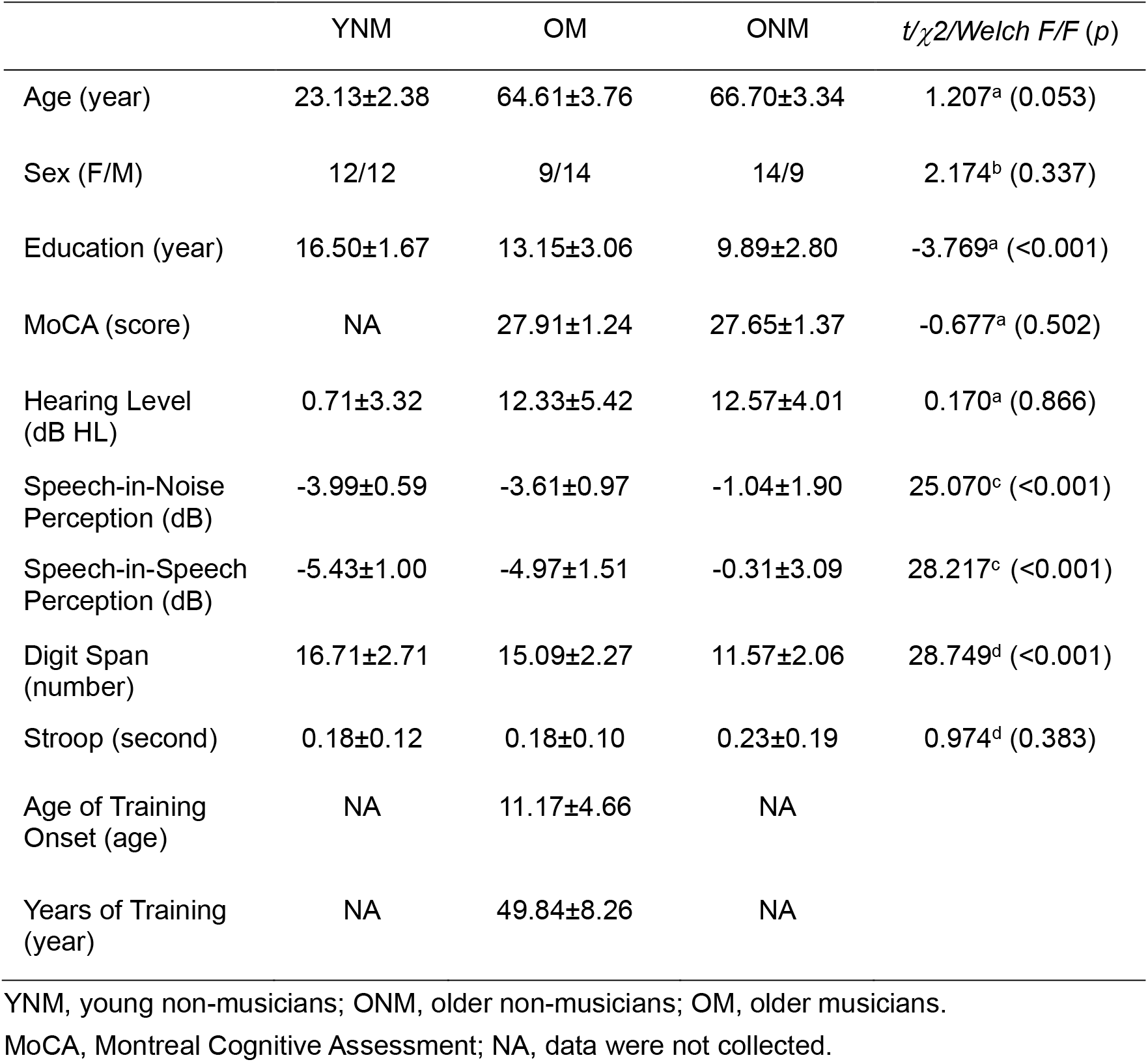
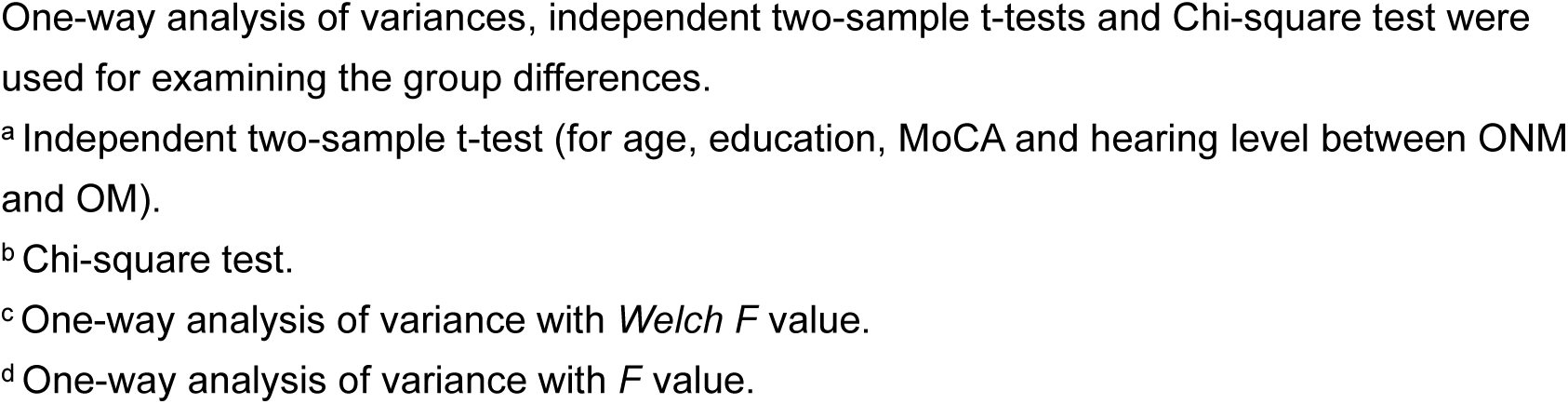
The group mean ± standard deviation values and statistics of demographic and behavioral data in each group.

| | YNM | OM | ONM | $t/\chi^2$/Welch $F/F$ ($p$) |
| --- | --- | --- | --- | --- |
| Age (year) | 23.13±2.38 | 64.61±3.76 | 66.70±3.34 | 1.207<sup>a</sup> (0.053) |
| Sex (F/M) | 12/12 | 9/14 | 14/9 | 2.174<sup>b</sup> (0.337) |
| Education (year) | 16.50±1.67 | 13.15±3.06 | 9.89±2.80 | -3.769<sup>a</sup> (<0.001) |
| MoCA (score) | NA | 27.91±1.24 | 27.65±1.37 | -0.677<sup>a</sup> (0.502) |
| Hearing Level (dB HL) | 0.71±3.32 | 12.33±5.42 | 12.57±4.01 | 0.170<sup>a</sup> (0.866) |
| Speech-in-Noise Perception (dB) | -3.99±0.59 | -3.61±0.97 | -1.04±1.90 | 25.070<sup>c</sup> (<0.001) |
| Speech-in-Speech Perception (dB) | -5.43±1.00 | -4.97±1.51 | -0.31±3.09 | 28.217<sup>c</sup> (<0.001) |
| Digit Span (number) | 16.71±2.71 | 15.09±2.27 | 11.57±2.06 | 28.749<sup>d</sup> (<0.001) |
| Stroop (second) | 0.18±0.12 | 0.18±0.10 | 0.23±0.19 | 0.974<sup>d</sup> (0.383) |
| Age of Training Onset (age) | NA | 11.17±4.66 | NA | |
| Years of Training (year) | NA | 49.84±8.26 | NA | |

YNM, young non-musicians; ONM, older non-musicians; OM, older musicians.
MoCA, Montreal Cognitive Assessment; NA, data were not collected.
One-way analysis of variances, independent two-sample t-tests and Chi-square test were used for examining the group differences.
<sup>a</sup> Independent two-sample t-test (for age, education, MoCA and hearing level between ONM and OM).
<sup>b</sup> Chi-square test.
<sup>c</sup> One-way analysis of variance with *Welch F* value.
<sup>d</sup> One-way analysis of variance with *F* value.

### Musical expertise offsets age-related hemispheric lateralization reduction

To reveal how aging and musical expertise influence brain functional lateralization, we examined the difference in LIs between YNM, ONM, and OM. We first calculated global LI maps of LI_intra and LI_he at the vertex level for each group, and found that YNM presented left-lateralized LI_intra in language related areas and DMN areas and right-lateralized LI_intra in CON areas (**Fig. 2A**), which were similar to previously reported patterns (33, 34). However, results of LI_he in three groups have not been reported in previous studies. Generally speaking, ONM showed more symmetrical FC pattern and substantially diminished lateralization pattern compared to YNM, while OM preserved more similar lateralization pattern to YNM. These differences in global lateralization pattern were further verified by the correlations between group-averaged LIs in YNM and those in two older groups across all vertices (**Fig. 2B**), with OM and YNM having higher *r* values than ONM and YNM (LI_intra: *z* = 15.608, *p* < 0.001; LI_he: *z* = 10.606, *p* < 0.001).

**Fig. 2.**
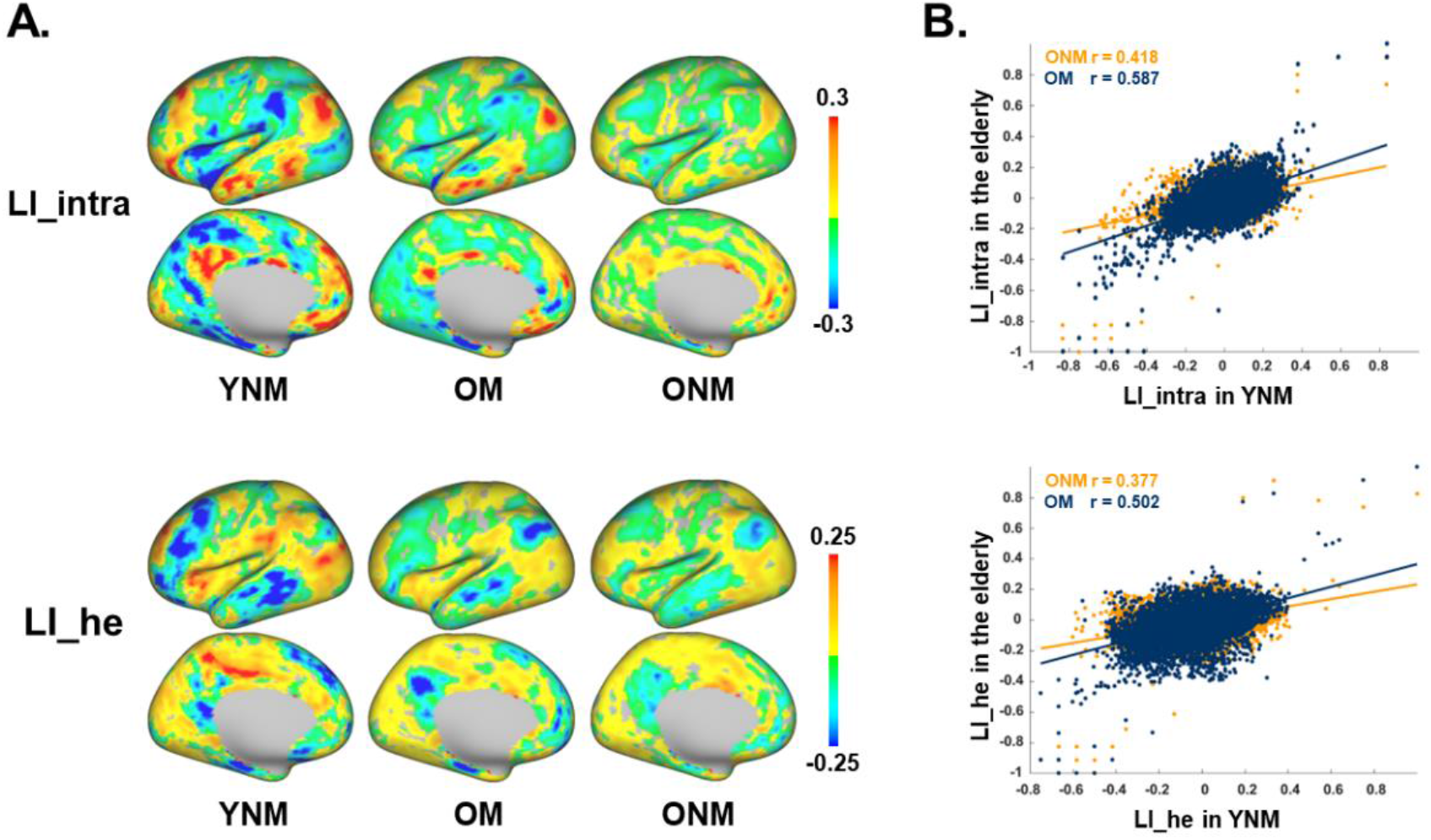
Vertex-based LI in three groups. (**A**) Group averaged global maps of vertex-based LI_intra and LI_he in YNM, OM, and ONM. (**B**) Correlations between group-averaged LIs in YNM and in two older groups across vertices, with OM showing significantly higher correlation than ONM (OM, blue; ONM, yellow).

To confirm our findings at the network level, we calculated network-based LI by averaging the LIs belong to the same network defined by Cole-Anticevic Brain-wide Network Partition version 1.0 (CAB-NP v1.0) (36). We selected seven networks that are relevant to speech in noise tasks (37) and modulated by aging and musical training (38), including somatomotor network (SMN), dorsal attention network (DAN), language network (LAN), auditory network (AUD), CON, FPN and DMN. We then defined network-based LIs based on global LI maps. One-way ANOVAs revealed a significant group difference on LI_intra of CON (*F*(2, 67) = 4.706, *η*^2^ = 0.123, *p* = 0.012) and LI_he of LAN (Welch *F*(2, 38.241) = 3.313, *η*^2^ = 0.093, *p* = 0.047). Post hoc tests demonstrated that ONM showed a significantly larger LI (close to zero) than YNM (LI_intra of CON: FDR-corrected *p* = 0.021; LI_he of LAN: Games-Howell-corrected *p* = 0.044) but no difference was discovered between OM and two non-musician groups (**Fig. 3**). Further one-sample t-tests found a significantly right-lateralized LI_intra of CON in YNM and OM (both *t* < –4.099, FDR-corrected *p* < 0.001) but not in ONM, and a significantly right-lateralized LI_he of LAN in all three groups (all *t* < –2.772, FDR-corrected *p* < 0.05). These results supported the HAROLD model and suggested that aging was associated with reduced hemispheric asymmetry especially in CON and LAN via functional compensation, while musical expertise counteracted such changes in lateralization via functional preservation, making OM more alike YNM.

**Fig. 3.**
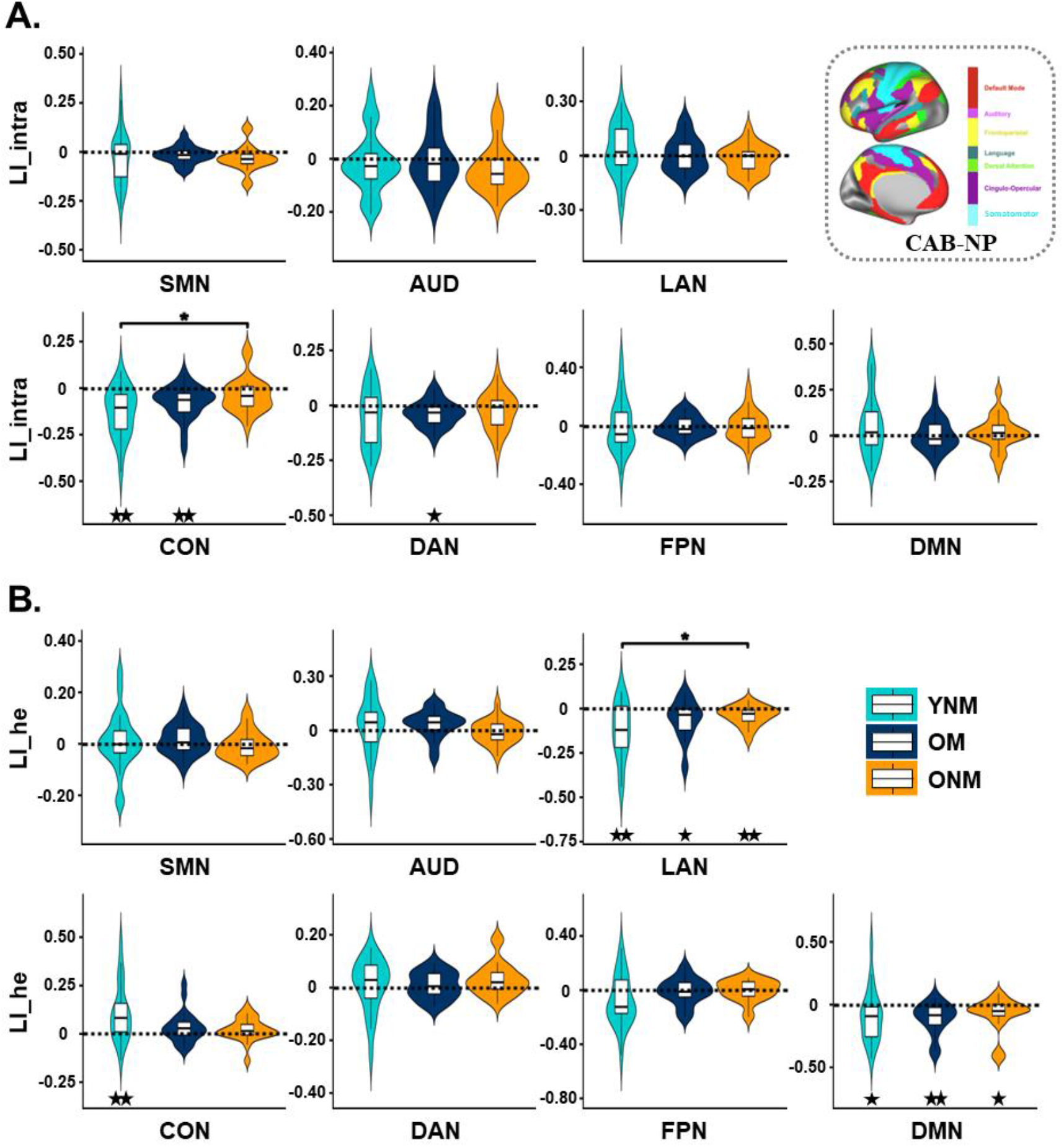
Group comparisons of network-based LI. LI_intra (**A**) and LI_he (**B**) were compared among YNM, OM and ONM in each of the seven networks. * FDR-corrected or Games-Howell-corrected *p* < 0.05 by post hoc tests after one-way analysis of variances. ^★^ FDR-corrected *p* < 0.05, ^★★^FDR-corrected *p* < 0.01 by one-sample t-tests. All dashed lines indicate zero (no functional lateralization).

### Musical expertise rejuvenates older adults’ lateralization pattern

As shown above, OM had more similar lateralization pattern as YNM. To directly verify this phenomenon, we adopted the neural alignment-to-young measure of network-based LI by calculating the cosine similarity between LI_intra/LI_he in every subject and the group average of LI_intra/LI_he in YNM across the vertices belonging to the same network (**Fig. 1C**). Lower neural alignment represents less similar functional lateralization pattern to YNM. One-way ANOVAs on neural alignment of LI_intra (**Fig. 4A**) demonstrated significant group differences in several networks including LAN, CON, FPN and DMN (all *F/Welch F* > 5.082, *η*^2^ > 0.135, *p* < 0.010). Post hoc tests revealed that ONM showed significantly lower alignment-to-young than YNM in all four networks (all FDR-or Games-Howell-corrected *p* < 0.01), but no significant difference was found between OM and YNM. In parallel, significant group differences were found on neural alignment of LI_he in SMN, CON, DAN, FPN and DMN (all *F* > 4.912, *η*^2^ > 0.128, *p* < 0.010, **Fig. 4B**). Post hoc tests showed significantly lower alignment in both ONM and OM than in YNM in SMN, CON, DAN and FPN (all FDR-corrected *p* < 0.05). For DMN, a significant difference was found only between YNM and ONM (FDR-corrected *p* = 0.014), but not between YNM and OM. Together, we confirmed that OM preserved youth-like lateralization pattern in networks including LAN, CON, FPN and DMN.

**Fig. 4.**
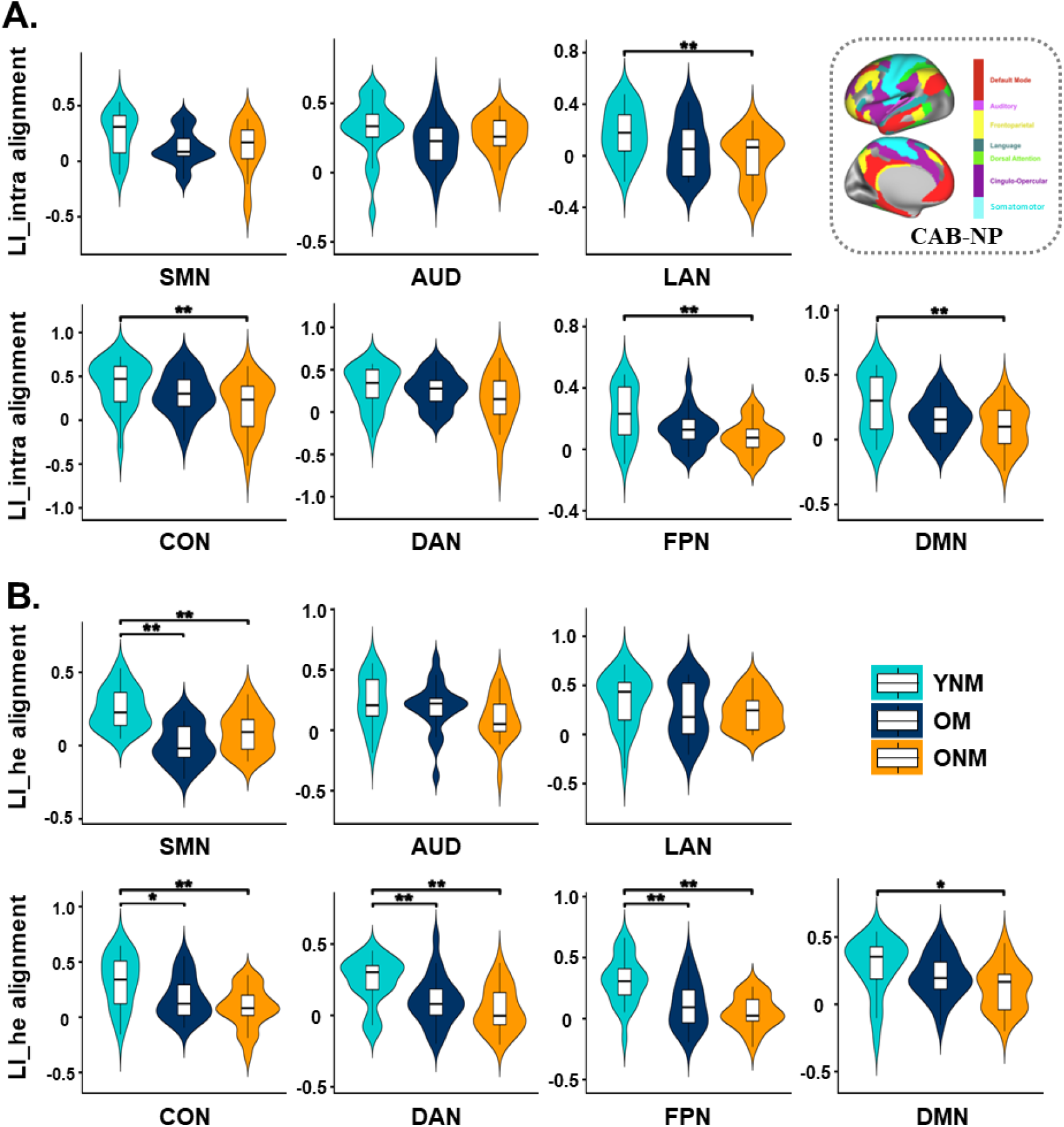
Group comparisons of neural alignment-to-young of network-based LI. Neural alignment-to-young of LI_intra (**A**) and LI_he (**B**) were compared among YNM, OM and ONM in each of the seven networks. * FDR-corrected *p* < 0.05, ** FDR-corrected or Games-Howell-corrected *p* < 0.01 by post hoc tests after one-way analysis of variances.

### Lateralization differentially contributes to speech perception in noisy environments in older musicians and older non-musicians

Since OM and ONM presented different lateralization patterns relative to YNM, we then asked whether and how functional lateralization was related to speech in noise perception performance in two older groups, of which brain network exhibiting functional compensation and which one producing functional preservation. Notably, hearing level, working memory and inhibitory control were related to speech perception in noise, and significant group differences were found in education, mean framewise displacement (mFD) (*t*_44_ = –2.147, Cohen‘s *d*= –0.633, *p* < 0.05). We thus regressed out sex, education, hearing level, digit span, stroop score, MoCA, mFD and mean global FC (*t*_44_ = –1.055, *p* = 0.297), and conducted partial correlations of network-based LI and its neural alignment with speech perception threshold in two older groups, respectively.

As shown in **Fig. 5A**, lower SIN or SIS threshold (representing better performance) was correlated with more left-lateralized LI_intra in SMN (SIN: *r* = –0.620, FDR-corrected *p* = 0.049) and DAN (SIN: *r* = –0.618, FDR-corrected *p* = 0.049; SIS: *r* = –0.517, uncorrected *p* = 0.048), and more left-lateralized LI_he in DMN (SIN: *r* = –0.554, uncorrected *p* = 0.032) in ONM. In contrast, in OM lower SIS threshold was correlated with more right-lateralized LI_intra in FPN (*r* = 0.690, FDR-corrected *p* = 0.031) and DAN (*r* = 0.617, uncorrected *p* = 0.014, FDR-corrected *p* = 0.050). Therefore, the two older groups displayed quite opposite relationships between functional lateralization and speech perception threshold.

**Fig. 5.**
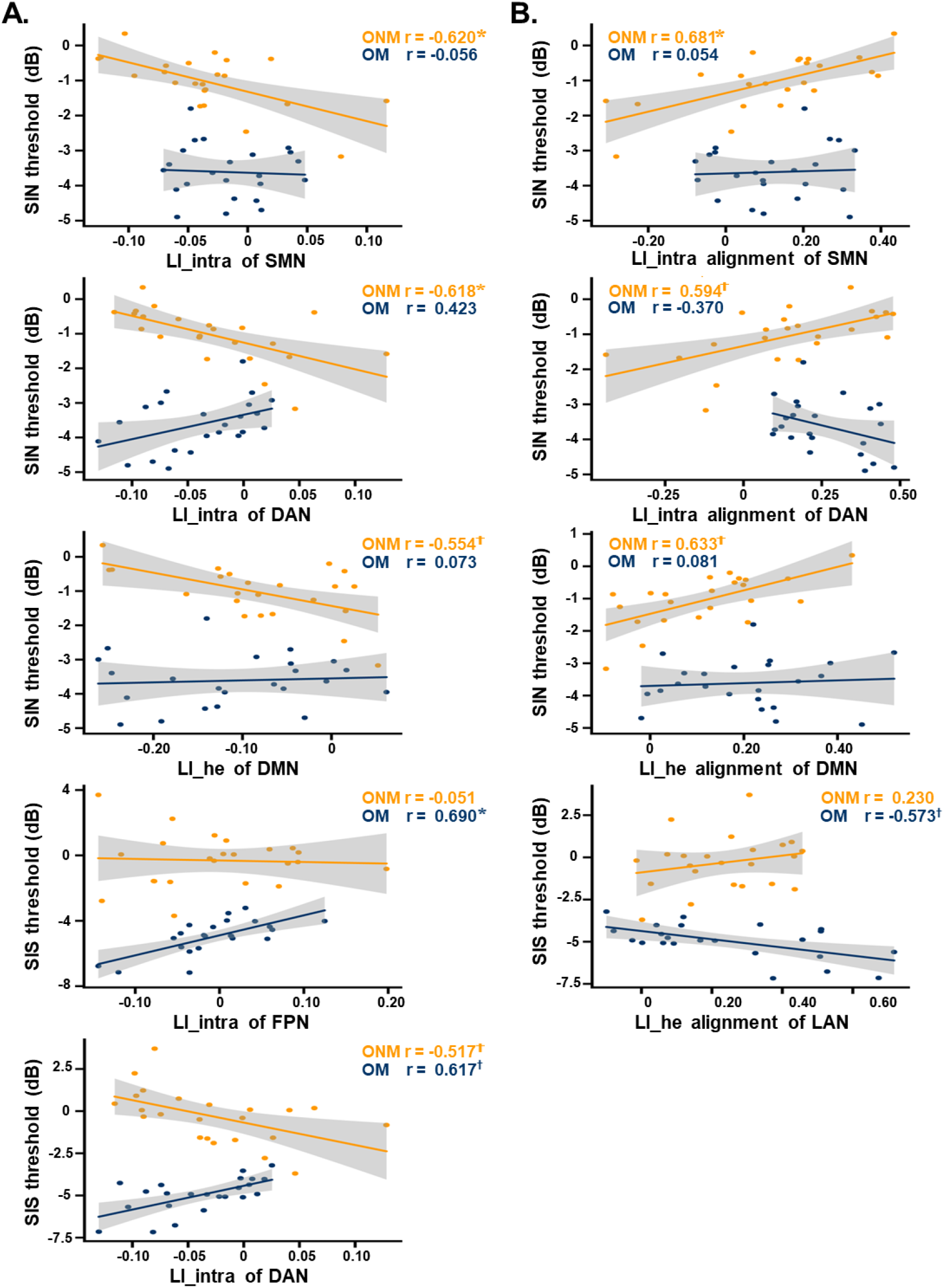
Correlations of network-based LI and its neural alignment with behaviors. (**A**) Partial correlations between network-based LI and speech-in-noise (SIN) threshold or speech-in-speech (SIS) threshold in OM and ONM, after controlling for sex, education, hearing level, digit span, stroop, MoCA, mean framewise displacement and mean global functional connectivity. (**B**) Similar partial correlations between network-based LI alignment and SIN or SIS threshold. * FDR-corrected *p* < 0.05, ^†^ uncorrected *p* < 0.05.

Furthermore, we investigated whether preserved speech in noise perception ability in older adults with or without lifetime musical experience was linked to a lateralization pattern similar to that of young adults or the opposite. By means of partial correlations between network-based LI neural alignment and speech perception threshold, in ONM lower neural alignment of LI_intra in SMN (*r* = 0.681, FDR-corrected *p* = 0.036) and DAN (*r* = 0.594, uncorrected *P* = 0.019) and lower alignment of LI_he in DMN (*r* = 0.633, uncorrected *p* = 0.011) were correlated with lower SIN threshold (**Fig. 5B**), that is, the less similar the lateralization was to YNM, the better performance, suggesting that functional compensation of lateralization may play an important role in speech perception in noisy conditions for older non-musicians. However, in OM higher alignment of LI_he in LAN (*r* = –0.573, uncorrected *p* = 0.026) was correlated with lower SIS threshold (**Fig. 5B**), that is, the more similar the lateralization was to YNM, the better performance, indicating an exactly opposite pattern of functional preservation of lateralization in speech perception in noisy conditions for older musicians.

## Discussion

From a previously untouched perspective of intrinsic brain functional lateralization, we provide evidence that long-term musical training counteracts age-related decline of speech in noise perception via preservation of youth-like functional lateralization, whereas older adults without musical training experience rely on stronger scaffolding of compensatory hemisphere with less similar functional lateralization pattern as young adults to maintain speech perception performance. In parallel with behavioral findings that age-related decline in perceiving speech sentences under “cocktail party” scenarios (with either speech or noise masker) was found only in ONM but not in OM, YNM’s strong right lateralization of intrahemispheric FC in CON and interhemispheric heterotopic FC in LAN were significantly decreased to a bilaterally symmetric pattern in ONM but not in OM. In addition, ONM were less similar to the YNM template than YNM in the functional lateralization of most networks, while OM preserved youth-like lateralization pattern in networks including LAN, CON, FPN and DMN. Moreover, functional lateralization contributed to speech in noise perception quite differently in OM and ONM: in ONM, the more left-lateralized pattern and the lower neural alignment-to-young (i.e., less similar lateralization pattern to YNM), the better performance, suggesting a functional compensation mechanism; whereas in OM, the more right-lateralized pattern and the higher neural alignment-to-young (i.e., more similar lateralization pattern to YNM), the better performance, suggesting a functional preservation mechanism. Therefore, successful aging in speech perception can be achieved by opposite functional lateralization changes in older adults with and without musical training.

### Functional compensation in older non-musicians

It is generally accepted that our brain responds to age-related anatomical and physiological changes by reorganizing its function (4). Compared with young adults, older adults showed a more bilateral pattern of prefrontal activity during verbal recall, which motivated the HAROLD model interpreting this functional lateralization change as reflecting functional compensation (4). Resting-state fMRI studies also reported lateralization decrease with aging in several networks, such as sensorimotor, attentional, and frontal networks (23, 39). All these findings convergently support the notion that the increased bilateral recruitment and lateralization decrease serve as a compensatory scaffolding for age-related neural decline (1).

Here, we found that LI_intra in CON and LI_he in LAN significantly changed from right-lateralization in YNM to bilateralization in ONM, which corresponded to previous studies and further clarified the specific need for additional neural resources for attentional control via left within-hemisphere interactions and language processing via left-to-right across-hemisphere interactions by older non-musicians. However, these LI changes for functional compensation did not directly influence speech in noise perception in ONM. Instead, ONM performed better with a less similar functional lateralization pattern to YNM presenting more left-lateralized in SMN, DAN, and DMN. Previous task fMRI studies have demonstrated the compensatory role of left speech motor areas (17) and bilateral sensorimotor regions (10) for poor speech encoding under adverse listening conditions in ONM. From the perspective of functional lateralization of resting-state fMRI, our findings that high-performing older adults had a more bilateral and even left-lateralized SMN which deviated from the pattern of young adults further supplemented the sensorimotor integration as a compensatory scaffolding for speech perception in the aging group. In addition, consistent with the posterior-anterior shift in aging and the decline-compensation hypothesis (40, 41), ONM also recruit more frontal areas to compensate for declined sensory functions when performing the SIN task (15–17). Since the ability to track and understand speech amid competing sound sources is supported by higher-level cognitive processes such as selective attention (42–44), the functional lateralization pattern of DAN and its relationship with speech in noise perception in high-performing ONM might represent another compensatory scaffolding for greater listening effort. Besides focusing attention on external auditory information, subjects need to inhibit task-unrelated long-term memory supported by DMN to avoid interference, which means that the DMN is required to be inhibited when perceiving speech in noise. Previous studies have reported older adults’ deficits in cognitive control and resource reallocation to the task-related regions, indicating that failure to inhibit DMN in the elderly is detrimental to task performance (45, 46). In high-performing ONM, we detected a less similar functional lateralization pattern to YNM displaying more bilateral and even leftward LI_he of DMN, which supported the compensation theory that higher-order cognitive networks may become more bilateral and even in the opposite lateralization pattern to compensate for sensory declines. In total, older adults without musical training experience rely on functional compensation of multiple networks controlling sensorimotor integration, attention and inhibition of long-term memory to maintain speech in noise perception performance.

### Functional preservation in older musicians

Broad factors (e.g., experience, genetics, and environment) are important determinants to influence the course of aging and, in turn, the level of cognitive function (5). Life-course variables, i.e., the accumulation of experiences and states an individual has experienced from birth to death (47), can impact the structure and function of the aging brain (5). Long-term musical training experience is one of those variables that have been found to improve auditory and cognitive functions during adverse listening environments, especially in the aging population (6, 9). Here, functional lateralization of CON and LAN in YNM was significantly weakened in ONM but was preserved in OM. Neural alignment analysis further detected more evident discrepancies between YNM and ONM than between YNM and OM in LAN, CON, FPN and DMN, demonstrating that musical training experience helps older adults preserve a youth-like functional lateralization pattern in networks association with language processing, attentional and cognitive control. These wide spread youthful lateralization patterns are consistent with previous researches (29, 33), showing that musical training has an age-decelerating effect on the brain (6, 9, 10). Compared to non-musicians, musicians’ predicted brain age was younger than their chronological age using a machine-learning algorithm (48).

Moreover, this functional preservation supported speech in noise perception in OM. A recent task fMRI study found that OM had better speech-in-noise perception performance through functional preservation by maintaining similar speech representation patterns as young adults (10). However, in this resting-state fMRI study, high-performing OM showed more rightward LI_intra in DAN and FPN and higher alignment-to-young of LI_he in LAN, presenting an exactly opposite relationship to ONM. LI_intra in DAN was significantly right-lateralized only in OM, while bilateral organized in YNM and ONM. These findings were consistent with previous studies revealing a bilateral dorsal attention system (49) and increased recruitment of the right hemisphere in speech processing in musicians (24, 25). Since speech in noise perception engages allocation of attentional resources and inhibitory control (50) that help to disentangle the target signal from the masker (51), enhancement of these higher-level cognitive processes has been discovered to correlate with improved speech in noise perception in musicians (9, 27, 52, 53). Due to the partial overlap between neural circuits dedicated to music and language (54), long-term musical training may influence language networks continuously and help preserve the youth-like lateralization pattern of LAN in older musicians. Our study supports this notion, as we found high-performing OM showed similar lateralization pattern with higher alignment-to-young of LI_he in LAN. The youth-like lateralization pattern of DAN, FPN and LAN might allow older musicians to functionally reserve cognitive abilities including language, selective attention, and inhibitory control that all contribute to better speech perception performance.

### Limitations

Since OM eligible and willing to participate in our fMRI study were rare, OM with different types of musical expertise were combined into one group (e.g., piano, singing, violin, etc.). However, the musical training type could have different effect on brain lateralization (55, 56). Comparing the effect of different types of lifelong musical training that older adults major in could further reveal how musical expertise promotes speech-in-noise perception via functional lateralization. In addition, the significant difference in neural alignment of network-based LI was only found between ONM and YNM, but not between ONM and OM, although OM did not differ from YNM in many networks. A larger sample of participants in future research will help verify the difference between OM and ONM directly.

To sum up, from a previously unidentified perspective of resting-state functional lateralization and its relationship with speech in noise perception, we found that 1) compared to young adults, older non-musicians show significant hemispheric lateralization reduction while older musicians keep youthful lateralization pattern; 2) to maintain speech in noise perception performance, older non-musicians rely on stronger scaffolding of compensatory networks with a more bilateral and even left-lateralized pattern, whereas older musicians depend on stronger preservation of youth-like functional lateralization with a more right-lateralized pattern. Thus, older non-musicians and older musicians form different coping strategies against aging, which helps deepen the understanding of functional compensation and functional preservation in aging theories and inspires individualized training intervention.

## Materials and Methods

### Participants

Seventy-four healthy native Mandarin speakers with no history of psychiatric or neurological disorders participated in the experiment, including 24 YNM (23.13 ± 2.38 years, twelve females), 25 ONM (66.64 ± 3.40 years, sixteen females) and 25 OM (65.12 ± 4.06 years, eleven females). All participants had normal hearing in both ears with average pure-tone threshold < 20 dB hearing level from 250 to 4,000 Hz. OM started training before 23 years old (mean = 10.90 ± 4.56 years old) with at least 32 years of training (mean = 50.88 ± 8.75 years), and practiced consistently in recent three years (1 to 42 hours per week, mean = 12.70 ± 8.99 hours per week). Non-musicians reported less than two years of musical training experience, which did not occur in the year before the experiment. To screen out people with mild cognitive impairment, all older participants passed the MoCA of the Beijing version (≥ 26 scores) (57). All participants reported their educational background and sighed informed written consent approved by the ethical committee of the Institute of Psychology, Chinese Academy of Sciences. For more details on participants, see Zhang et al. (10) which used the same subjects.

## Behavioral Tests

The speech in noise perception threshold was assessed using the SIN and SIS tasks in which syntactically correct but semantically meaningless speech sentences (for instance, “一些条令已经翻译我的大衣” “Some rules had translated my coat”) were embedded in a speech-spectrum noise (SIN task) or a two-talker speech masker (SIS task) at different signal-to-noise ratios (SNR = −12, −8, −4, 0, and 4 dB). The stimuli were presented binaurally through Sennheiser HD380 Pro headphones driven by a Dell desktop computer. The interaural time difference of the target sentence and the masker was manipulated to generate two perceived spatial relationships between the target and the masker: colocation and separation. Participants were asked to repeat the whole target sentence as best as they could immediately after the sentence was completed. A logistic psychometric function was employed in Matlab 2016b to fit each subject’s data for each masking and spatial relationship condition using the Levenberg– Marquardt method, and the SNR corresponding to 50% correct identification across two spatial relationships was used as the threshold ratio for the SIN and SIS tasks. For more details on other tests we used, see the corresponding section in Supporting Information.

### Data Acquisition and Processing

Imaging data were collected using a 3 T MRI system (Siemens Magnetom Trio) with a 20-channel head coil. The high-resolution T1-weighted anatomical image was obtained using magnetization-prepared rapid acquisition with gradient echo (MPRAGE): repetition time (TR) = 2200 ms, echo time (TE) = 3.49 ms, field of view (FOV) = 256 mm, flip angle (FA) = 8°, slice thickness = 1 mm, voxel size = 1 × 1 × 1 mm, 192 slices. Slowly fluctuating brain activity was measured using a multiband-accelerated echo-planar imaging (EPI) series with whole-brain coverage while subjects were instructed to rest still and quietly: TR = 640 ms, TE = 30 ms, FOV = 192 mm, FA = 25°, slice thickness = 3 mm, voxel size = 3 × 3 × 3 mm, 40 slices, multiband factor = 4. Each T1-weighted scan lasted 7 min 13 s and each resting-state scan lasted 8 min 6 s for a total of 750 consecutive whole-brain volumes.

Preprocessing was performed using fMRIPrep 20.2.5 (58). Functional MRI data were preprocessed with slice-timing correction, motion correction, distortion correction, co-registration to structural data, normalization to MNI space, and projection to cortical surface. Then the eXtensible Connectivity Pipeline (XCP-D) (59) was used to post-process the outputs of fMRIPrep, including demeaning, detrending, nuisance regression, band-pass filtering. In addition, we used FD to identify high movement frames in data (>0.5 mm). By adopting “scrubbing” for each of these data points (head radius =50 mm for computing FD, FD threshold = 0.5 mm for censoring), we excluded two OM with more than 20% data above the high motion cutoff (FD > 0.5). Moreover, one ONM for lacking complete behavioral data and one ONM who was left-handed were also excluded. Therefore, seventy right-handed subjects were included in the subsequent analysis. For more details on data processing, see the corresponding section in Supporting Information.

### Definition of LI

To quantify rs-FC at the surface level, we downsampled the original 32k time series into those with 10k vertices to accelerate further steps. As typically, the rs-FC between two cortical surface vertices was computed as Pearson correlation (r) of two vertex-wise blood-oxygen-level dependent (BOLD) time series and then converted Fisher’s r-to-z transformation to improve the normality.

Based on the whole-brain FC matrix, we obtained interhemispheric heterotopic FC and intrahemispheric FC. The former represented the FC between two cortical surface vertices across different hemispheres, except the homotopic pairs while the latter indicated the FC between two cortical surface vertices within the same hemisphere. For a specific cortical surface vertex, the heterotopic (he) was defined as the sum of heterotopic FCs between this vertex and all the others in the opposite hemisphere except the homotopic one, whereas the intrahemispheric (intra) was defined as the sum of intrahemispheric FCs between this vertex and all the others within the same hemisphere. On these bases, we further defined two different forms of functional lateralization, calculated as

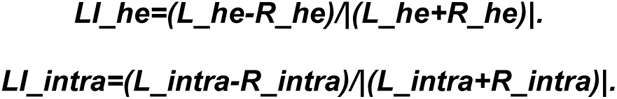

Larger positive values of LI_he and LI_intra imply stronger bilateral across-hemisphere interactions or ipsilateral within-hemisphere interactions in left-hemispheric vertices, whereas larger negative values indicate stronger interactions in right-hemispheric vertices. Finally, every subject had two LI maps of LI_he and LI_intra.

### Definition of Network-based LI and Its Neural Alignment

According to the CAB-NP v1.0 (36), all cortical surface vertices were mapped into twelve networks. To calculate network-based LI, we only chose the homotopic pair of vertices belonging to the same networks via downsampled CAB-NP v1.0 from 32k to 10k. After averaging within the same network, seven network-based LIs relevant to our behavior tasks were acquired, including SMN, AUD, LAN, CON, DAN, FPN, and DMN. We further employed an inter-subject pattern correlation framework to examine spatial network-based LI similarities between every subject and the YNM average (10, 60). For each network-based LI in every subject, we adopted an alignment-to-young measure by directly comparing the LI pattern of each subject and the mean LI pattern of YNM across the same network, using cosine similarity. By doing this, three cosine similarity matrices of YNM, ONM, and OM for seven networks were obtained. Higher neural alignment implies more similar functional lateralization pattern to YNM, whereas lower neural alignment indicates less similar functional lateralization pattern to YNM. All analyses above were performed on both LI_he and LI_intra.

### Statistical Analysis

To highlight the differences in LI_he and LI_intra between the three groups, we presented the averaged global maps of LIs in YNM, ONM, and OM, respectively. Next, we performed one-sample t-tests on seven network-based LIs separately for each group to identify whether this functional network was lateralized (leftward/rightward) or symmetric. One-way ANOVAs for network-based LIs and network-based LI neural alignments were adopted to compare the group differences.

Since we intended to explore the relation between functional lateralization and behavior, partial correlations were used to evaluate the relationship between network-based LIs as well as their neural alignments and speech in noise tasks in two older groups, considering sex, education, mFD, mFC (mean of rs-FCs between any two cortical surface vertices in the whole brain), hearing level, auditory digit span, stroop, and MOCA (older subjects only) as confounding variables. All the results in our analyses were considered to be significant with the value below 0.05 after FDR correction or Games-Howell correction (one-way ANOVA with heterogeneity of variance). The brain maps were projected to surfaces depicted by SurfStat package (www.math.mcgill.ca/keith/surfstat) and Connectome Workbench software platform (http://www.humanconnectome.org/software/connectome-workbench.html).

## Acknowledgments

The authors thank Yining Chen and Yiyang Wu for helping with data collection.

## Funding

STI 2030—Major Projects 2021ZD0201500 (YD)

National Natural Science Foundation of China 31822024 and 31671172 (YD)

Strategic Priority Research Program of Chinese Academy of Sciences XDB32010300 (YD)

Scientific Foundation of Institute of Psychology, Chinese Academy of Sciences E1CX172005 (XHJ), E2CX3625CX (YD) and E1CX4725CX (XYW)

## Data and materials availability

The data that support the findings of this study are available in https://osf.io/msz2r/.

