## Supporting Information for "Compensation or preservation? Different roles of functional lateralization in speech perception in older non-musicians and musicians"

**Behavioral Tests.**

*Digital span task*

Auditory working memory was measured using the forward and backward Digit Span subtest of the Wechsler Adult Intelligence Scale of Chinese version (1). Participants were presented with a recording of a series of digits spoken by a young Chinese female. The number of digits increased from 3 to 12 in the forward part and from 2 to 10 in the backward part (two trials per length), while participants were asked to repeat the digits in a normal and reverse order, respectively. The task stopped if both trials for the same length were incorrect. Auditory working memory score was defined as the sum of the longest numbers participants could repeat in the forward and backward parts. Note that, raw scores instead of age-normed scores were used for analysis because the age difference was the research interest in the current study. Larger values indicate better performance.

*Stroop task*

The Stroop test was adopted to evaluate inhibition control ability (2). Participants were asked to listen to the Chinese words then press the response key in synchronous condition (e.g., “high (高)” in high pitch, the lexical meaning and pitch were always congruent), and in asynchronous condition (e.g., “high (高)” in low pitch, the lexical meaning and pitch were always incongruent). Performance was indexed as the following equation: time of asynchronous condition minus time of synchronous condition. Larger values indicate poorer performance.

**Data Processing.**

Preprocessing was performed using fMRIPrep 20.2.5 (3); RRID:SCR_016216), which is based on Nipype 1.6.1 (4); (5); RRID:SCR_002502). Structural data were corrected for intensity non-uniformity, skull-stripped, and then used for reconstruction of the cortical surface. Volume-based structural images were segmented into cerebrospinal fluid (CSF), white matter, and gray matter, then spatially normalized to the standard MNI space. Functional MRI data were preprocessed with slice-timing correction, motion correction, distortion correction, co-registration to structural data, normalization to MNI space, and projection to cortical surface. Functional timeseries were resampled to FreeSurfer’s fsaverage space, and grayordinates files containing 91k samples were generated.

Built with Nipype 1.7.0 (4), the eXtensible Connectivity Pipeline (XCP-D) (6) was used to post-process the outputs of fMRIPrep. For each CIFTI run per subject, all data were subjected to demeaning, detrending, and nuisance regression. Volumes with framewise displacement (FD) greater than 0.5 mm (7, 8) were flagged as outlier and excluded from nuisance regression. The nuisance regression pipeline was based on the empirical tests performed by Ciric et al. 2017 (9). Specifically, six primary motion parameters were removed, along with their derivatives and the quadratics of all regressors (24 motion regressors in total). Physiological noise was modeled based on white matter and ventricle signals using aCompCor (10) within fMRIprep. Five component signals were used, as well as their derivatives and the quadratics of all physiological noise regressors (20 physiological noise regressors in total) (11). Next, the residual time series from this regression were then band-pass filtered to retain signals within the 0.01-0.08 Hz frequency band. After smoothed with a gaussian kernel size of 6 mm (FWHM) using FMRIB Software Library (FSL), processed functional time series were extracted from residual blood-oxygen-level dependent (BOLD) using Connectome Workbench (12). For more details, see the xcp_d website (<https://xcp-d.readthedocs.io>).

1.	Y. Gong, Manual of Wechsler adult intelligence scale-Chinese version. *Changsha: Chinese Map* (1992).
